# Simulations and Electrostatic Analysis Suggest an Active Role for DNA Conformational Changes During Genome Packaging by Bacteriophages

**DOI:** 10.1101/288415

**Authors:** Kim A. Sharp, Xiang-Jun Lu, Gino Cingolani, Stephen C. Harvey

## Abstract

Motors that move DNA, or that move along DNA, play essential roles in DNA replication, transcription, recombination, and chromosome segregation. The mechanisms by which these DNA translocases operate remain largely unknown. Some double-stranded DNA (dsDNA) viruses use an ATP-dependent motor to drive DNA into preformed capsids. These include several human pathogens, as well as dsDNA bacteriophages – viruses that infect bacteria. We previously proposed that DNA is not a passive substrate of bacteriophage packaging motors but is, instead, an active component of the machinery. Computational studies on dsDNA in the channel of viral portal proteins reported here reveal DNA conformational changes consistent with that hypothesis. dsDNA becomes longer (“stretched”) in regions of high negative electrostatic potential, and shorter (“scrunched”) in regions of high positive potential. These results suggest a mechanism that couples the energy released by ATP hydrolysis to DNA translocation: The chemical cycle of ATP binding, hydrolysis and product release drives a cycle of protein conformational changes. This produces changes in the electrostatic potential in the channel through the portal, and these drive cyclic changes in the length of dsDNA. The DNA motions are captured by a coordinated protein-DNA grip-and-release cycle to produce DNA translocation. In short, the ATPase, portal and dsDNA work synergistically to promote genome packaging.

## Introduction

DNA translocases – motors that move DNA, or that move along DNA – are required for numerous processes during the cell cycle. A key question is that of energy transduction: what is the mechanism by which the energy released by the biochemical cycle (ATP binding and hydrolysis, and product release) is converted to the mechanical cycle of DNA motion? The packaging motors of double-stranded DNA (dsDNA) bacteriophages are among the strongest of all biological motors (1, 2), and they are ideal systems for investigating energy transduction in one class of translocases.

dsDNA viruses infect a wide range of organisms. Among these are the herpesviruses, which infect almost every tissue type in the human body. They are responsible for a number of diseases, including chicken pox and shingles; infectious mononucleosis; retinitis pigmentosa; and oral and genital lesions. The tailed bacteriophages (order *Caudovirales*) provide simple model systems for herpesviruses. Like them, the phages have dsDNA genomes, but unlike them, the phages are unenveloped. The packaging motors of herpesviruses and other dsDNA pathogenic viruses are potential targets for novel antiviral agents. Our research is aimed at elucidating the mechanism by which the motor drives dsDNA into the capsid.

Assembly of tailed phages begins with the spontaneous formation of a procapsid whose structure is based on icosahedral geometry. At one vertex of the procapsid there is an ATP-dependent motor that drives the dsDNA genome into the empty procapsid. The motors of the three phages studied here – φ29, T4 and P22 – contain two protein components, a pentameric ATPase and a dodecameric portal. (The nomenclature in this field is not uniform; the φ29 portal is commonly called the “connector”, and the ATPases of some of these viruses – but not φ29 – are called "terminases". Some authors use "motor" to indicate the ATPase, but there is growing evidence that conformational changes in the portal are part of the packaging mechanism (3, 4), so we use “motor” to indicate the ATPase-portal complex.) The φ29 motor is unusual in that it also includes a pentameric RNA component (pRNA). More detail on dsDNA bacteriophages and their motors is available in several excellent reviews (5–10).

In the mature virus, the DNA is packed quite densely. As a consequence, the motor must overcome strong forces arising from the electrostatic DNA-DNA repulsions (11–16), the cost of bending the DNA double helix (11–17), and the entropic penalty arising from the drastic reduction in the conformational space available to the DNA (15, 16, 18). With regard to the mechanism of energy transduction by phage motors, the details of the biochemical cycle remain poorly understood, but a number of models have been put forward for the mechanical cycle (19–33). Almost all of these assume that the DNA is a passive substrate, and that a protein-DNA grip-and-release cycle is coupled to protein motions that push (or pull) the DNA along.

The first suggestion that the DNA might play an active role in packaging was Lindsay Black’s “torsional compression translocation” model (19), in which DNA is “crunched” by the motor proteins, storing elastic energy that is later released to move the DNA forward. The model is supported by evidence for DNA shortening during packaging in phage T4, including changes in fluorescence resonance energy transfer (FRET) (21) and the ejection of intercalating agents as the DNA is packaged (34). This model invokes lever-like motions in the ATPase to generate “a linear force that locally compresses duplex DNA”, *e.g.*, Fig. 7 of (19) and Fig. 4 of (21).

We suggested an alternate hypothesis for the cause of DNA shortening. In this "scrunchworm model" (31), the proteins cycle between conformations that alternately dehydrate and rehydrate the DNA. This produces a cycle of transitions between the B-DNA and A-DNA conformations. A-DNA is shorter than B-DNA, and coupling these transitions to a coordinated grip-and-release cycle rectifies the DNA motions. There is no intermediate storage of elastic energy in the DNA. Our molecular dynamics (MD) simulations on DNA in the channel of the φ29 connector showed that the DNA is shortened, or "scrunched" (35). The same conformation was observed in simulations on the same system by the Grubmüller lab (36). Neither of these simulations included lever-like protein motions, so they both supported the scrunchworm model.

This paper describes the results of computer simulations on dsDNA in the channels of the φ29, T4 and P22 portals. In the simulations, the length of the DNA double helix depends on the protein sequence, and, for P22, on the conformation of the portal. Further analysis reveals that the DNA responds to the electrostatic potentials generated by the portals. This suggests a mechanism by which the energy released by the chemical cycle is harnessed to drive the DNA into the capsid.

## Objectives and Approach

Our long-range goal is to examine the interactions between DNA and the phage motor. There are no high-resolution structures for any of the motors, so we began our studies by examining the behavior of dsDNA in the channel of four structures for the portal assemblies, one each from φ29 and T4, and two different conformations of the P22 portal.

We hypothesize that the range of differences in these four structures resembles the range of conformational changes of a single portal as it passes through the steps of the chemo-mechanical cycle. If this is true, there might be significant differences in the optimal (lowest energy) DNA conformations in these four complexes. We tested this idea by introducing a segment of double-stranded DNA into the channel of each of the portals, running MD simulations on the portal-DNA complex while restraining the portal, and examining the DNA conformations after they reached equilibrium.

Our study was aimed at answering four questions: Are the optimal DNA structures in all of these portals similar? If not, what are the characteristics of these conformations? Are the DNA conformations unusual, or do they resemble conformations previously described in the literature? And finally, by what mechanism does the portal structure drive the differences in DNA conformation?

## Results

### DNA assumes a portal-dependent conformation

Fig. 1 shows the conformation of the DNA when equilibrated against four different portal structures, those of φ29, T4, and two different conformations of the P22 portal. One of these (“P22-MV”) is from the 3.25Å crystal structure of the dodecameric portal bound to 12 copies of gp4 (37); this structure has 12-fold symmetry. The other (“P22-PC”) is from the 3.3Å crystal structure of the isolated portal core, which is asymmetric (4). The nomenclature reflects the hypothesis that the portal is asymmetric in the procapsid (PC) but is driven to the symmetric conformation upon DNA packaging, yielding the mature virion (MV) (4).

**Fig. 1.**
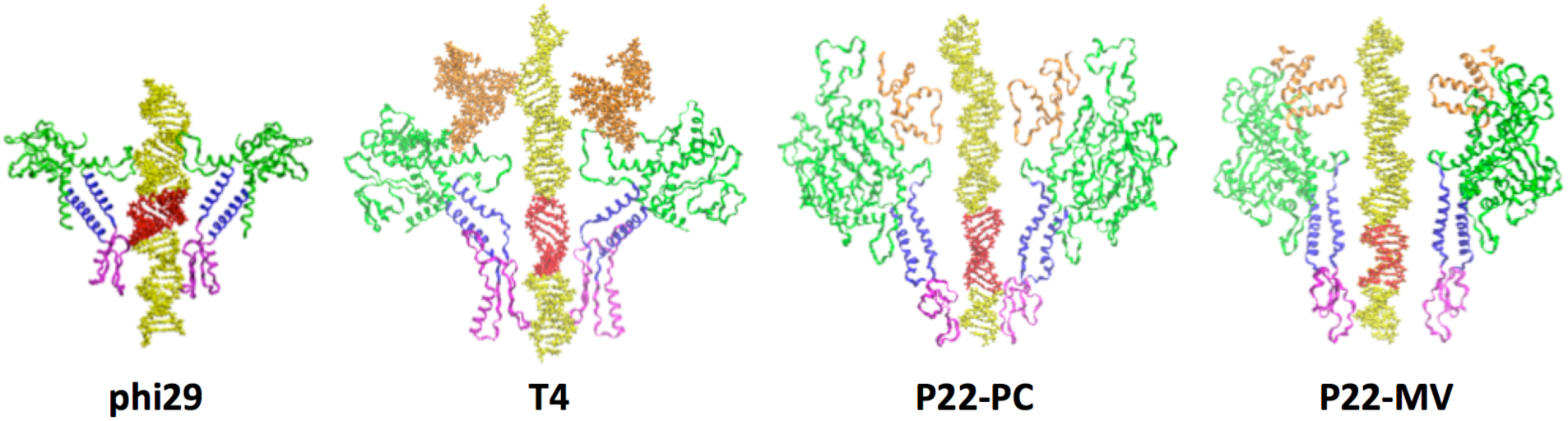
The equilibrium conformation of DNA in the viral portals depends on the sequence and structure of the portal protein. These cross-sectional views show two of the protein chains from opposite sides of the dodecameric portal. Protein domains are colored using the scheme introduced by Sun *et al.* (39): green, blue, magenta and orange identify the wing, stem, clip, and crown domains, respectively. The principal variations in DNA conformation (red) occur near the bottom of the alpha-helical stem region. Note that the P22 models include only the core region of the protein, because the barrel domain is not resolved in the PC structure (4), and it was determined only at low resolution in the MV structure (37).

The equilibrium DNA conformations are quite varied in the four portal structures, particularly in the region near the base of the alpha helical "stem" domain of the portal (Fig. 1, red base pairs). The differences are manifested by differences in the pitch of the double helix or, equivalently, by the rise per base pair.

As we previously reported (35), a segment of DNA in the φ29 portal is "scrunched", with an average rise of 2.1 ±0.2 Å; this is shorter than B-DNA (rise 3.4Å) and even shorter than A-DNA (rise 2.6Å). The scrunching is independent of DNA sequence, as it occurred in DNAs containing four different sequences with a wide range of A-phobicities (35). ("A-phobicity" is the resistance to the B-to-A transition in the presence of dehydrating agents (38)). Kumar and Grubmüller reported a similar structure in their simulations on the same complex (36).

As seen in Fig. 1, DNA lengthens in the T4 portal, and in the P22 portal in the PC conformation; in each of these, there are several base pair steps with helix rise of 4Å or more. Interestingly, DNA in the MV conformation of the P22 portal is neither shortened nor lengthened, remaining close to the B-DNA conformation.

The most significant result is that different conformations of the same viral portal produce different DNA conformations (P22-MV *vs.* P22-PC). This supports the scrunchworm hypothesis that the cycle of ATP binding, hydrolysis and product release causes protein conformational changes that drive shortening and lengthening motions of the DNA (31). It does, however, refute the proposal that DNA cycles between the A- and B-forms (31).

It is interesting that the DNA conformational changes all occur in the same region of the channel, but it is puzzling that the DNA is shortened in one case, and that it is lengthened in two cases. This naturally leads to the question: what drives these protein-dependent differences in DNA conformation?

### Protein-DNA contacts do not explain the changes in DNA conformation

Fig. 2 shows all the proteins atoms that are within 5Å of the DNA in each of the four portal-DNA complexes. In the φ29 portal, as we previously reported (31), there are several protein contacts with the backbone of the scrunched DNA, particularly with the "load-bearing" strand (40). In contrast, there are almost no protein-DNA contacts in the two complexes where the DNA is extended (T4 and P22-PC), or in the P22-MV complex, where the DNA is essentially B-form.

**Fig. 2.**
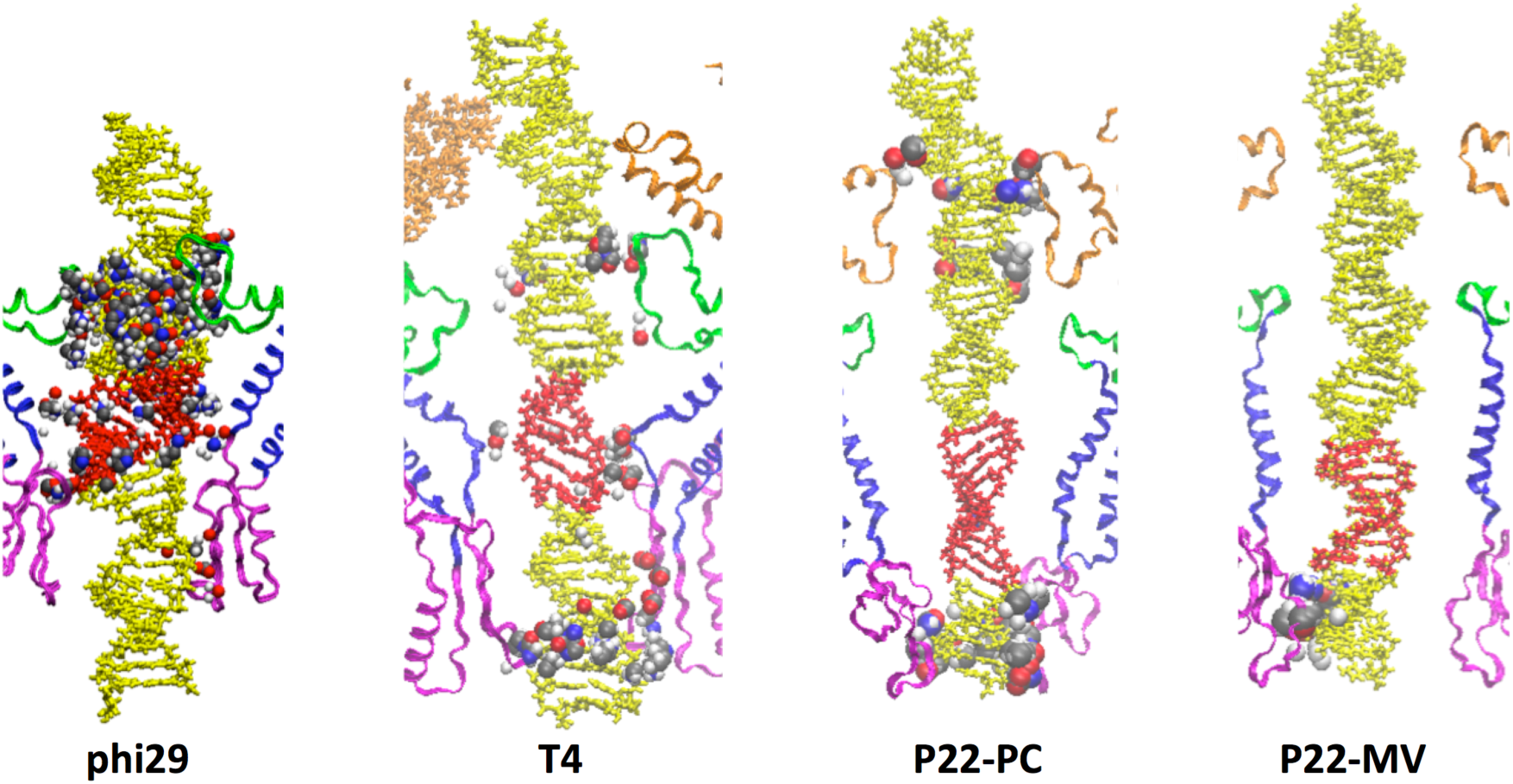
Protein-DNA contacts. Grey, red, blue and white spheres represent atoms of carbon, oxygen, nitrogen and hydrogen, respectively, within 5Å of any DNA atom for the four structures shown in Fig. 1. The scrunched region in phi29 has several contacts, particularly along one backbone, as described previously (35). In contrast, the stretched regions in the T4 and P22-PC DNAs have almost no direct contacts. Most important, the difference between the DNA conformations in P22-PC and P22-MV DNAs is clearly not due to direct contacts with the portal protein.

### DNA conformation is driven by the electrostatic potential pattern in the channel

Fig. 3 shows the electrostatic potential maps for the four portal structures, and they explain the differences in DNA conformation seen in Fig. 1. The negatively charged phosphate groups are drawn into regions of significant positive potential (>*k_B_T*/*e*) and repelled from regions of significant negative potential (<–*k_B_T*/*e*). (*k_B_* is Boltzmann’s constant, *T* is the absolute temperature, and *e* is the charge on the proton.) Consequently, DNA is shortened in regions of positive potential and lengthened in regions of negative potential.

**Fig. 3.**
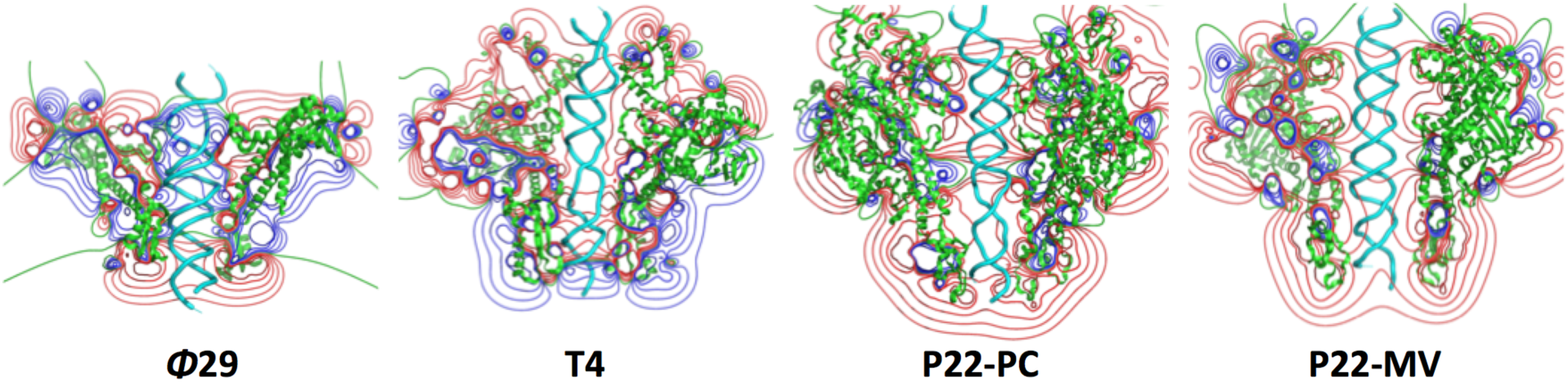
Electrostatic potential maps in the channels of the four portals shown in Fig. 1 (cross section). Positive (blue) and negative (red) contours are shown for ±0.25, ±0.5, ±1, ±2 and ±4 *k_B_T*/*e*. In φ29, the phosphates are drawn toward a ring of positive charges lining the channel, causing the DNA to shorten. In T4 and P22-PC, the phosphates are repelled from regions of high negative potential, causing DNA lengthening. The electrostatic potential in the P22-MV channel is apparently not large enough to drive a significant conformational change in the DNA.

The inward compression of the phosphate groups in regions of high negative potential pushes them away from the channel walls. This is why there are few contacts between the portal proteins and the stretched DNA conformations in T4 and P22-PC (Fig. 2).

These results argue that cyclic conformational changes in the viral motor could produce an electrostatically-driven cycle of DNA shortening and lengthening. The question then becomes, how to rectify these DNA conformational changes to produce translocation? The answer suggests a novel motor mechanism, as described below (Discussion).

### In regions with large positive base pair inclination, the sign of base pair slide determines whether DNA is shortened or lengthened

The distorted DNA conformations (red in Fig. 1) are associated with runs of large positive base pair inclination. Shortening happens with runs of negative base pair slide (A-DNA and φ29), while stretching occurs with runs of positive slide (>2Å in T4 and P22-PC). This agrees with the observation of Lu and Olson (41) that DNA is stretched if slide and inclination have the same sign, and it is compressed when they have opposite signs. Fig. S1 (see Supporting Information) is a scatter plot of inclination and slide, showing the role of slide-inclination correlations in determining DNA compression and elongation.

### Structural homology searches show that scrunched DNA structures are common, but the stretched DNA structure in the T4 and P22-PC simulations is new

We used two approaches to search through all DNA structures in the Protein Database (PDB) (42), to see if similar DNA conformations exist in isolated dsDNA molecules, or in protein-DNA complexes. In the first search, each structure was scored using the root mean square difference (RMSD) between atoms in the target structure and corresponding atoms in the structure being queried. The second search used base pair slide as a query, since A-DNA and the scrunched φ29 DNA have large negative slide, while the stretched DNA segments in the T4 and P22-PC simulations have large positive slide.

By either criterion, there are many DNAs in the PDB database with conformations similar to the scrunched DNA in the φ29 channel, which resembles A-DNA. The results using the RMSD criterion are tabulated in Table S1 (Supporting Information). The closest hit (PDB ID 3J9X) is the DNA of the rod-shaped SIRV2 virus, all of which is covered by protein, driving it to the A-form (46). It is shown in Figure S2. The abundance of A-DNA-like structures in DNA-protein complexes has been reported before (43). This is not surprising, since protein binding often strips solvent off the DNA, and dehydration has long been known to drive DNA toward the A-form (44, 45). DNA-RNA complexes generally have the A-form, too, and there are several examples of this in Table S1.

Using our first search criterion (lowest RMSDs), there are a number of stretched DNA conformations similar to those seen in our simulations. The P22-PC search required a match of 7-10 base pairs, and it produced only nine hits with RMSDs < 3Å (Table S2 and Figure S3 in Supporting Information). The strongest hits are in complexes of DNA with bound intercalating drugs, other ligands, or covalent modifications. There are only two complexes of a protein bound to an unmodified DNA. These have RMSDs of 2.9Å from the target DNA structure, barely meeting the 3Å cutoff. The T4 search used a shorter template, requiring a match over 5-6 base pairs. There were 74 hits with RMSD < 3Å (Table S3). Again, the large majority of the complexes (54/74) involve bound intercalating drugs, other ligands, or covalent DNA modifications. The closest hits for protein-DNA complexes are the same as those found in the P22-PC search. Visual inspection and analysis of the remaining protein-DNA complexes by curves+ and 3DNA show that the DNAs do not have the large values of rise, positive slide and positive inclination characteristic of the stretched DNAs in the simulations.

Our second search sought structures with positive base pair slide, averaging more than 2.0Å over at least five successive base pairs, to match the stretched regions in the T4 and P22-PC structures. This turned up only one stretched structure (PDB ID 3VH0). This is a complex between a protein and an 11-nucleotide single-stranded DNA; the structure contains a DNA duplex due only to crystallographic symmetry, and it is a badly distorted duplex. There is little base pairing, and two bases are bulged outside the duplex. Total helix rise is only 24Å over eight bp steps, similar to B-DNA (27Å). This is much less than the rise for eight base pairs of stretched DNA in our simulations (37Å for P22-PC, and 36Å for T4). In short, 3VH0 bears no resemblance to the stretched DNA in the P22-PC and T4 simulations, a result again confirmed by visual inspection.

The combined results of our database searches suggest that the stretched structures in our P22-PC and T4 simulations represent a previously unobserved DNA conformation.

## Discussion

Our simulations show that phosphate groups are drawn together in regions of high positive potential, causing DNA shortening, and phosphates are pushed apart in regions of high negative potential, causing DNA stretching. This suggests that DNA is actively involved in force generation and translocation, as shown schematically in Fig. 4.

**Fig. 4.**
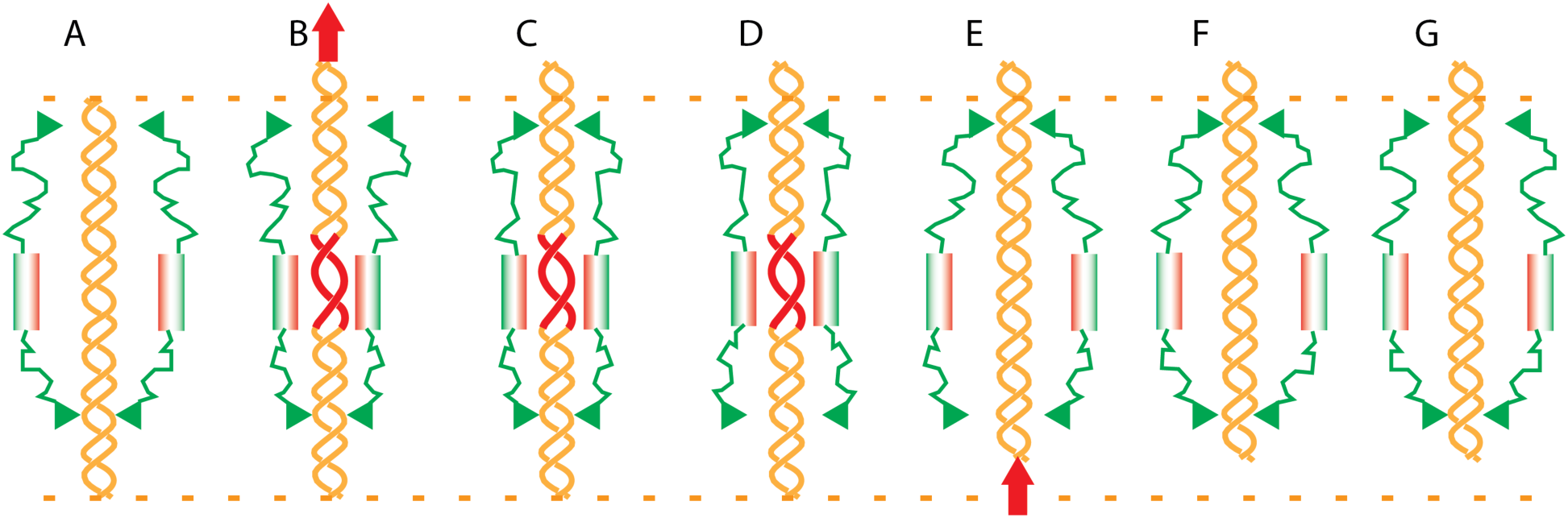
DNA translocation can be driven by coupling cyclic changes in electrostatic potential with cyclic motions of two protein-DNA grips. (A) B-DNA (orange) in the channel of the complex of motor proteins (two protein chains shown schematically in green). The lower protein-DNA grip is closed at the bottom of the channel, and the upper grip is open (pairs of green triangles). (B) A conformational change in the proteins brings a negatively charged protein domain (pink) close to the DNA, driving the nearby DNA into an extended form (red). The head of the DNA is pushed forward into the capsid (arrow). (C) The upper grip closes to capture the DNA’s advance. (D) The lower grip opens. (E) A second conformational change in the protein pulls the negatively charged domain away from the DNA. The DNA returns to the B-form, pulling the tail of the DNA upward. (arrow). (F) The lower grip closes to prevent DNA backsliding. (G) The upper grip opens, returning the protein to its original conformation. The DNA has advanced upward from its original position, which is marked by the dashed orange lines.

We propose that the energy released by the biochemical cycle drives cyclic conformational changes in the motor proteins, moving a charged protein domain close to the DNA at one part of the cycle (Fig. 4A), and further away later in the cycle (Fig. 4E). This causes changes in the electrostatic potential acting on the DNA, which drives a DNA lengthening-shortening cycle. DNA motions are rectified by coordinating the changes in electrostatic potential with cyclic opening and closing motions of protein-DNA grips. Grips have been invoked in every model for the translocation mechanism (19–31).The same mechanism can be used if the protein charges are positive and DNA alternates between B-DNA and a compressed conformation. In this case, DNA shortening would draw the tail of the molecule into the channel, and the return to the B-form would propel the head of the molecule out of the channel and into the capsid.

At present, it is not possible to specify exactly how different conformations of the portal proteins would produce the systematic changes in the electrostatic potential described by the model. But the portal is a dodecamer, so electrostatic effects of even modest movements of the charged groups can be amplified by as much as twelve-fold. The charge motions might be produced by changes in the portal’s symmetry, as happens in the transitions between the P22-MV and P22-PC conformations. Alternatively, the alpha helices in the stem region might rotate around their long axes, or those axes might change their polar and azimuthal angles relative to the *z*-axis. (Note the differences between the orientations of the stem helices in the MV and PC conformations in Fig. 1.) Such motions were proposed in the first report of a portal’s crystal structure (47). It is also currently unclear which regions of the motor proteins would provide the grips postulated by the model, although the channels of all the portals studied here are narrow at both the upper and lower ends (Fig. 1). It is also possible that the upper grip is, in fact, a "one-way valve", as proposed by Guo and colleagues (32, 33).

We hypothesize that changes in the pattern in electrostatic potential are a key component of the mechanism connecting the energy released by ATP hydrolysis to DNA translocation. But it is important to acknowledge that we have not proven this. At present, there are no structures of the full complex between a viral ATPase and a portal, so we cannot yet define the successive conformational changes generated by the chemical cycle. Nor can we map the electrostatic potential in the channel of the complete ATPase-portal complex.

It remains possible that the ATPase is, as often assumed, the part of the motor that engages the DNA and drives it forward. If so, this suggests an alternative role for the electrostatic effects described here: they might be the basis for a protein-DNA grip within the portal channel. DNA movement would be hindered if the DNA contains a tract with a scrunched or stretched conformation like those seen in the φ29, T4 and P22-PC channels in Fig. 1. To move in either direction, the DNA would have to convert from the B-form to the scrunched (or stretched) conformation as it enters the tract, and it would have to make the reverse transition as it leaves the tract. Thus, if DNA axial motions are produced by the ATPase, electrostatic DNA-portal interactions could provide the grip required to rectify the motion. Once the structure of the full motor is known, simulations and electrostatic analyses can clarify the probable roles of individual charged amino acids, and those predictions can be tested by assays on engineered motors with suitable point mutations.

The φ29 motor is "promiscuous", in that it is able to package dsDNA substrates with short tracts containing a variety of modifications (40). The authors of that study argued that this is due to extensive protein-DNA contacts. Our φ29 simulations (35) revealed a number of non-specific protein-DNA contacts, and the protein-phosphate contacts were predominantly on the strand packaged in the 5’-to-3’ direction, which had been experimentally identified as the "load-bearing" strand (40). We cannot, however, fully explain the promiscuity of the φ29 motor, because the structure of the ATPase is unknown, and there are no data on the interaction of the DNA substrate with the ATPase. Resolution of this issue will require computer simulations on substrates in a complete motor (a complex of the dodecameric portal with the pentameric ATPase). Unlike the DNA shortening seen in φ29, the T4 and P22 motors cause DNA elongation, and there are almost no direct protein-DNA contacts in these two motors (Fig. 2). As a consequence, our model predicts that T4 and P22 will be much more sensitive to phosphate methylation – which neutralizes the charge – than is φ29. In support of this, the sensitivity of T4 to DNA modifications has some differences from that of φ29 (19).

Taken together, these results suggest that, when it comes to DNA translocases, "motor" is not always synonymous with "protein". In the viruses studied here, DNA is probably an active component of the motor, not a passive substrate, as traditionally assumed. If electrostatic interactions do connect the chemical cycle to DNA translocation in the manner proposed here, then there are two major issues to be resolved. First, there are not yet sufficient data to define the interactions between the DNA, the ATPase and the portal at each step of the packaging cycle. Second, It remains to be determined how the events of the biochemical cycle drive successive conformational changes in the proteins that, in turn, drive the DNA elongation/shortening cycle. Finally, if this model is substantiated by future research, it will be interesting to examine the possibility that a similar mechanism might be used by other translocases.

## Methods

### Portal structures

The 2.1Å crystal structure of the φ29 portal (PDB ID 1HSW) (48) is missing residues A230-S244. Kumar and Grubmüller modeled this segment and added it to the crystal structure, producing a model of the complete dodecamer (49), We used their model in our φ29 simulations, as described previously (35).

Our simulations on the T4 portal used the 3.6Å structure from cryo-electron microscopy (PDB ID 3JA7) (39).

We used two published structures for the P22 portal. One of these is a new refinement of the portal in the "MV" conformation (PDB ID 5JJ3) (4), based on the 12-fold symmetric 3.25Å crystal structure of the dodecameric portal protein associated with twelve copies of the P22 gene product 4 tail accessory factor (PDB ID 3LJ4) (4, 37); it includes residues 5-463 and 493-602. We also examined the 3.3Å crystal structure of a second conformation, designated "PC" (PDB ID 5JJ1) (4). The P22 portal has an extended barrel above the crown domain, but we omitted it, because it was seen only at low resolution in the MV conformation, and it was not seen in the PC structure.

### DNA models

We generated all-atom DNA models using the Web 3DNA server (50) (w3dna.rutgers.edu). We simulated four different sequences of dsDNA in the channel of the φ29 connector, as reported previously (35). We simulated the T4 channel with a 35 bp dsDNA (GAGATAACGATGCTCCACGAACTCCGGATTATCGG) and used a 40 bp dsDNA (GAGAGATAACGATGCTCCACGAACTCCGGATTATCGGGTG) for both P22-PC and P22-MV. Note that the central 35 bp of this molecule have the same sequence as the DNA in the T4 simulations. In each case, the 5’ end of the strand whose sequence is given is at the upper end of the portal, using the orientation in Fig. 1.

**MD simulations** were carried out in a solution of 150 mM NaCl in TIP3P water (51), using NAMD (52), the CHARMM36 force field (53, 54), a 2 fs time step, SHAKE restraints on bonds (55), particle mesh Ewald (PME) (56, 57) and the isobaric-isothermal (NPT) ensemble with rectangular repeating boundary conditions. Protein and DNA were restrained during 4 ns, for solvent equilibration. Production runs, with the protein restrained, ranged from 10-50 ns, with the aim of determining equilibrium DNA conformations driven by defined protein conformations. Structural convergence was determined by visual inspection, and by monitoring DNA helicoidal parameters and the A-B Index (58). As reported previously (35), the DNA equilibrated very rapidly – typically 1-2 ns – presumably because of the very strong electrostatic forces.

**Electrostatic potential maps** were calculated using DelPhi (59, 60).

### Structural homology searches

To compare the distorted dsDNA structures observed in the simulations with known DNA structures, we examined all DNA-containing entries in the RCSB PDB database (www.rcsb.org) (42) (6011 entries as of June, 2017). We used two search criteria. First, we calculated the RMSD between the atoms in a template DNA fragment from our simulations (scrunched or stretched) and the corresponding atoms in target dsDNA fragments from the database, using a sliding window. Bases were replaced by pseudoatoms defined by the standard base reference frame (61), to make the search independent of sequence. The closest hits were ranked by RMSD. The results are tabulated in Tables S1-S3 (Supporting Information).

In the second search, we calculated the slide and rise between every pair of contiguous base pairs in each structure of the database, using the standard definitions (61, 62) and an in-house implementation of the Curves+ algorithm (63, 64). Only hydrogen-bonded, base-paired double-stranded regions, with no bulged bases, were considered.

## Foot notes

Author contributions: S.C.H. and K.A.S. designed the research. S.C.H. carried out the simulations on DNA in the portals of φ29, T4 and P22. K.A.S. did the Delphi calculations and developed and applied the program for identifying structural homologues based on the slide criterion. X.J.-L. developed and applied the program for identifying structural homologues based on the RMSD criterion and contributed analyses using 3DNA. G.C. provided the structure of the P22-PC portal and insights on the differences between the PC and MV conformations. All authors participated in the discussions leading to the conclusions reported here, and in writing the manuscript.

The authors declare no conflict of interest.

## Acknowledgments

We are grateful to Rajendra Kumar and Helmut Grubmüller for providing us with the coordinates of their model for the φ29 portal, J.C. Gumbart for advice on the MD simulations, and Venigalla B. Rao for candid discussions. Supported by grants from the National Institutes of Health (R01 GM096889 to X-JL and R01 GM100888 to GC). MD simulations were carried out on the TACC Stampede system at the University of Texas and the Bridges system at the Pittsburgh Supercomputing Center; those resources are part of the Extreme Science and Engineering Discovery Environment (XSEDE), supported by National Science Foundation Grant ACI-1548562.

